# indCAPS: A tool for designing screening primers for CRISPR/Cas9 mutagenesis events

**DOI:** 10.1101/196121

**Authors:** Charles Hodgens, Zachary L. Nimchuk, Joseph J. Kieber

## Abstract

Genetic manipulation of organisms using CRISPR/Cas9 technology generally produces small insertions/deletions (indels) that can be difficult to detect. Here, we describe a technique to easily and rapidly identify such indels. Sequence-identified mutations that alter a restriction enzyme recognition site can be easily distinguished from wild-type alleles using a cleaved amplified polymorphic sequence (CAPS) technique. If a restriction site is created or altered by the mutation such that only one allele contains the restriction site, a polymerase chain reaction (PCR) followed by a restriction digest can be used to distinguish the two alleles. However, in the case of most CRISPR-induced alleles, no such restriction sites are present in the target sequences. In this case, a derived CAPS (dCAPS) approach can be used in which mismatches are purposefully introduced in the oligonucleotide primers to create a restriction site in one, but not both, of the amplified templates. Web-based tools exist to aid dCAPS primer design, but when supplied sequences that include indels, the current tools often fail to suggest appropriate primers. Here, we report the development of a Python-based, species-agnostic web tool, called indCAPS, suitable for the design of PCR primers used in dCAPS assays that is compatible with indels. This tool should have wide utility for screening editing events following CRISPR/Cas9 mutagenesis as well as for identifying specific editing events in a pool of CRISPR-mediated mutagenesis events. This tool was field-tested in a CRISPR mutagenesis experiment targeting a cytokinin receptor (*AHK3*) in *Arabidopsis thaliana*. The tool suggested primers that successfully distinguished between wild-type and edited alleles of a target locus and facilitated the isolation of two novel *ahk3* null alleles. Users can access indCAPS and design PCR primers to employ dCAPS to identify CRISPR/Cas9 alleles at http://indcaps.kieber.cloudapps.unc.edu/.

## Introduction

It is often necessary to genotype biological samples to select individuals from a large population with a desired genetic variant. Genetic variants generated by mutagenesis or natural variation can take the form of single nucleotide polymorphisms (SNPs) or insertions/deletions (indels). Sufficiently large indels can be distinguished using PCR followed by agarose or polyacrylamide gel electrophoresis (PAGE), but differences of one or two base pairs can be difficult to distinguish reliably even with PAGE, and SNP alleles are refractory to size-based genotyping. Diagnostic tools for genotyping samples with SNPs or small indels include PCR-based cleaved amplified polymorphic sequences (CAPS) or derived CAPS markers (dCAPS) [1]. A typical CAPS assay consists of a short amplicon centered on a restriction site present in only one genotype. The CAPs assay identifies the genotype of the individual based on whether or not the PCR product is cleaved by the differential restriction enzyme (Fig 1A). A dCAPS assay can be used if there are no restriction sites differentially present in the wild-type and mutant genomic sequences.The dCAPS assay introduces or disrupts a restriction enzyme motif near the mutation by amplifying the target sequences using an oligonucleotide primer that includes one or more mismatches relative to the template (Fig 1B). The mismatches are chosen so that following amplification, a restriction site is introduced into either the wild-type or the mutant amplified fragment, which can then be distinguished by restriction enzyme digestion followed by agarose gel electrophoresis.

**Fig 1.**
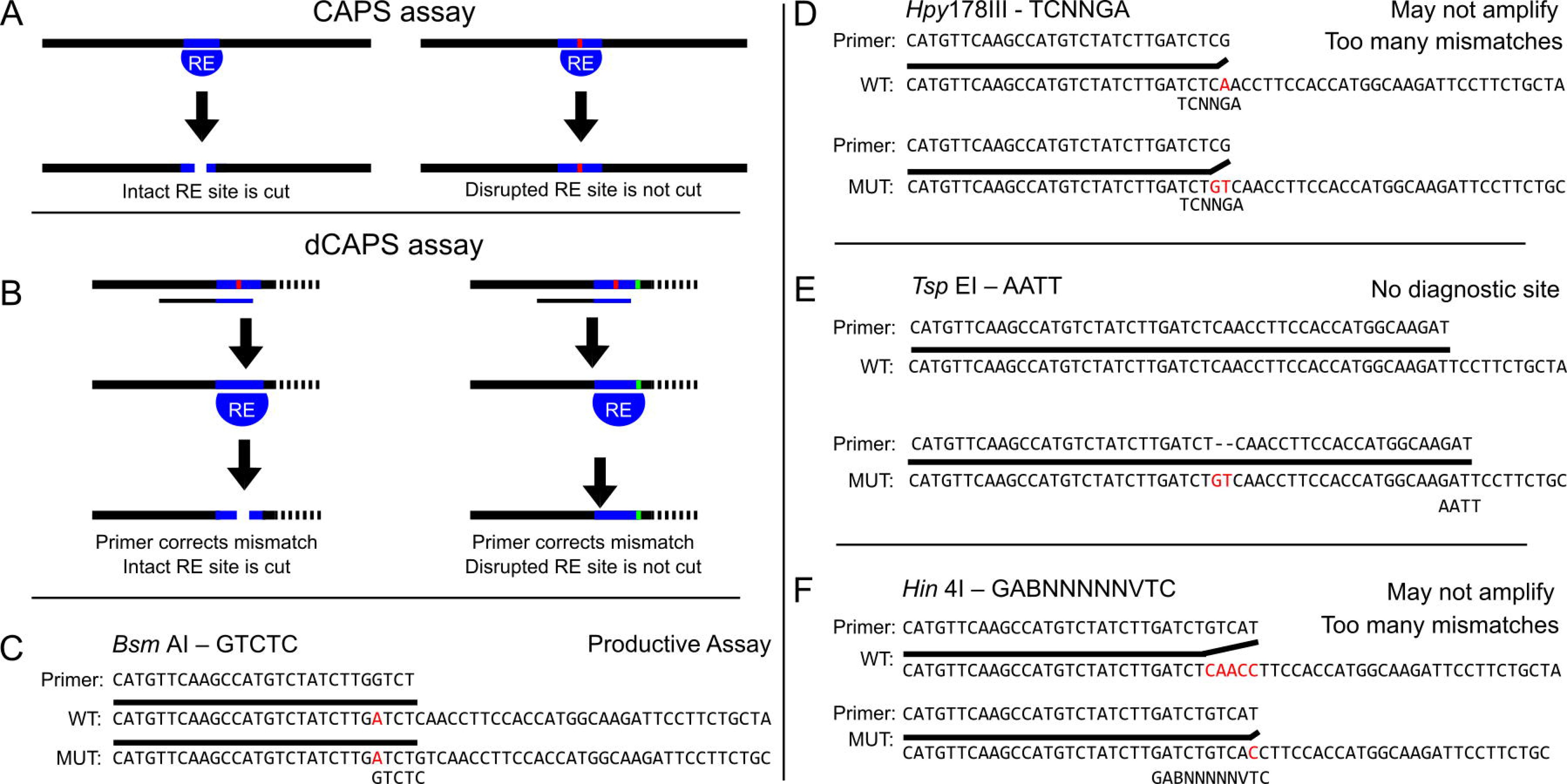
CAPS/dCAPS markers can distinguish alleles, but output of dCAPS Finder 2.0 can be flawed. (A) Diagram of CAPS technique. An amplicon centered on a restriction site (blue bar) disrupted by a SNP or indel (red bar) is differentially cleaved by a restriction enzyme (RE) in the wild-type vs mutant. (B) Diagram of the dCAPS technique. A restriction site can be introduced into either the wild-type or mutant target sequences using mismatched oligonucleotide primers to discriminate two sequences. The mutation (green bar) disrupts the introduced restriction site such that it is not cleaved by the restriction enzyme (RE). Gel electrophoresis can be used to identify the size difference between the wild-type and mutant fragments in both the CAPS and dCAPS methods. (C-F) A sequence with a two base pair deletion at a CRISPR cut site, chosen using CRISPR-Plant [8], was supplied to dCAPS Finder 2.0 with a mismatch allowance of 1 base pair.A minority of proposed assays are viable (C), but others possess too many mismatches for successful amplification by PCR or do not introduce diagnostic restriction sites (D-F).

As CRISPR/Cas9-generated mutant alleles become more prevalent, there is a growing need for a facile method for screening and genotyping indel alleles. Assays based on dCAPS markers are ideal for this as they are simple, robust, inexpensive, and relatively high throughput. However, designing productive primers for allele-specific dCAPS assays can be cumbersome.

Here, we present the development of a new web-based tool to design dCAPS primers for indels that should be of general utility for analysis of CRISPR/Cas9-generated mutant alleles in any species. We demonstrate the utility of this tool using CRISPR/Cas9 targeting of the *AHK3* locus in *Arabidopsis thaliana*. AHK2, AHK3, and AHK4 are the three receptors present in Arabidopsis that are involved in the perception of cytokinin, a plant hormone regulating a diverse set of biological functions in plants [2]. Previous studies have identified null alleles of *ahk2* and *ahk4*, but the previously identified *ahk3-3* allele is hypomorphic rather than a null allele, as residual full-length *AHK3* transcript was found to be present in seedlings harboring the strongest *ahk3* allele [3]. Primers generated by the indCAPS tool were successful at identifying editing events at the *AHK3* locus, and viable triple null mutant lines for the cytokinin histidine kinase receptors were identified. The indCAPS tool has the potential to be an important resource for investigators seeking to find new CRISPR alleles or design genotyping primers for known alleles.

## Materials and Methods

### Software

The indCAPS package was written in Python (version 3.5.2) and is implemented as a web application using the flask framework (0.12), the bleach package (1.5.0) for input scrubbing, and the gunicorn WSGI HTTP server (19.7.1). It is provided through an OpenShift application platform available from UNC-Chapel Hill. The website is available at http://indcaps.kieber.cloudapps.unc.edu. The source code is available at https://github.com/KieberLab/indCAPS.

### Plant Growth and Transformation

Plants were grown at 21°C in long days (16 h light). The *ahk2-7 cre1-12* double mutant was transformed with pCH59, a pCUT series binary expression vector containing *AHK3*-targeting gRNA sequences and expressing plant codon optimized Cas9 [4], by the floral dip method [5]. Putative transformants were selected on Murashige and Skoog media containing 50μg/ml hygromycin and then transferred to soil and allowed to set seed. T2 seeds were plated on Murashige and Skoog plant growth media (2400 mg/L MS salts, 250 mg/L MES buffer) [6]. Plants with long roots were transplanted to soil and genotyped for editing at the *AHK3* locus.

### Detection of *ahk3* mutations using dCAPS

Oligonucleotide primers were designed to detect editing at the *AHK3* locus using the indCAPS tool. Amplification of the *AHK3* locus was performed with primers AHK3.dC.F and AHK3.dC.rc (Table 1) followed by digestion of the amplicon with *Bsa* BI. Digests were analyzed using gel electrophoresis with a 3% agarose gel. Lines lacking any wild-type digestion pattern were selected for analysis. Sanger sequencing was used to characterize editing events as single base-pair indels and to confirm homozygosity.

**Table 1:**
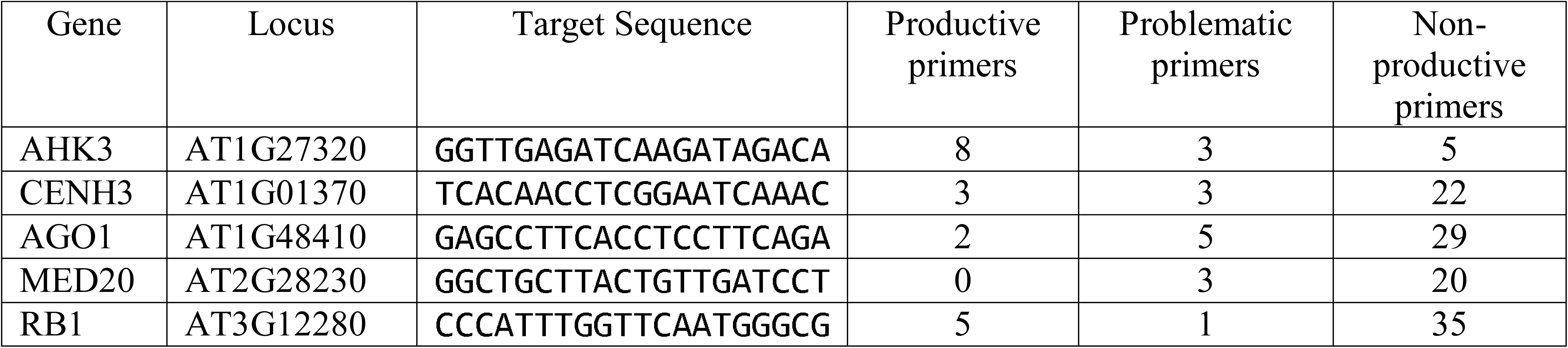
**Number of productive primers generated for tested loci.** Simulated amplicons were made using generated primers. Non-productive primers did not amplify sequences capable of being distinguished with a restriction digest. Problematic primers may not amplify due to features like 3’ mismatches or gaps in alignment to provided sequence.

**Table 2:**
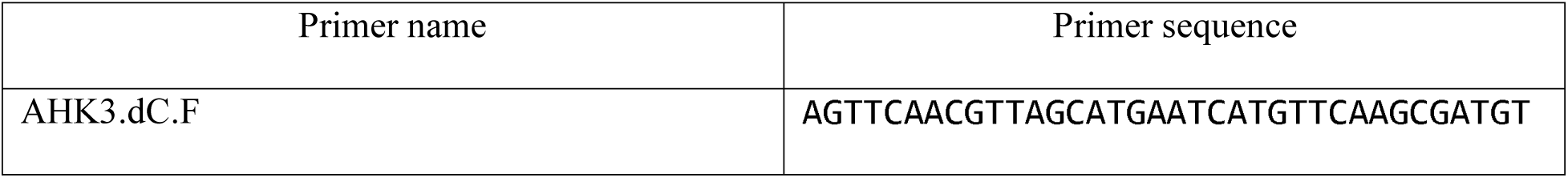
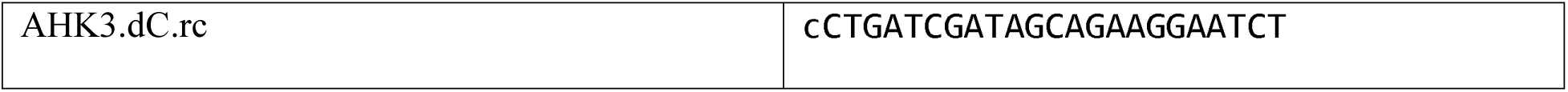
Primers used for CRISPR screening

## Results and Discussion

### dCAPS Finder 2.0 has poor compatibility with indel alleles

While a web tool for the design of dCAPS primers has been described [7], it was designed primarily to detect SNP alleles. Primers generated with the tool for small indels often will not actually amplify either the wild-type or mutant sequences by PCR, or in some cases will not actually distinguish between the wild-type and mutant sequences. For example, an analysis of potential dCAPS primers generated by the existing dCAPS program (http://helix.wustl.edu/dcaps/dcaps.html) for indels in several genes in *A. thaliana* demonstrated that as few as 15% of the suggested primers are capable of distinguishing the provided alleles (AHK3, 50% capable; CENH3, 14%; AGO1, 17%; MED20, 13%; RB1, 15%). The reason dCAPS Finder 2.0 falters on indel alleles is not clear as the source code is no longer available. The non-functional primers either do not generate a diagnostic restriction site or likely would not amplify the target DNA due to alignment gaps or extensive 3’ mismatches between the primer and one of the template sequences (Fig 1C-F). There are workaround methods in which the user supplies two sequences in which the terminal base is the indel, rather than placing the indel in the middle of the provided sequences. This approach will ignore any potential assays in which a restriction motif may overlap the indel site, as the program has no information about bases on the other side of the indel.

### A web-based tool for design of primers to detect indels: indCAPS

A new software package, indCAPS, was developed to facilitate the design of dCAPS primers for indels. This software has also been adapted for the design of CAPS and dCAPS oligonucleotide primers used in PCR amplification of target sequences in order to screen for editing events following CRISPR/Cas9-mediated mutagenesis [9]. The tool is available at http://indcaps.kieber.cloudapps.unc.edu.

The interface presents two dialog boxes to the user. The first box is for the generation of dCAPS primers for known alleles. The second box is for the generation of dCAPS primers for detecting unknown alleles. The first box requires the user to submit two sequences. No assumptions are made about either sequence being a wild-type or mutant allele, so order does not matter. Ideally, each sequence is centered on the mutation of interest. The two sequences do not need to be the same length, but should have homology arms of at least 20 bases flanking the mutation of interest. The user is also asked to submit a maximum number of mismatches in the primer. The default value is 1 mismatch. Increasing the maximum mismatch value should result in more enzymes being reported, but as with any hand-made dCAPS assay, this may result in primers which are less likely to successfully amplify DNA.

Several advanced options are also available. Amplicon length can be specified. This parameter dictates how far downstream the tool examines the sequence when looking for exact matches for each restriction site. A restriction enzyme is rejected if there is a cleavage site in the shared downstream region, which would complicate the analysis of the diagnostic cleavage at the site of interest. Depending on the size of the submitted sequence, the amplicon length may be longer than the sequence available to the program. In this case, the entire submitted sequence downstream of the primer is considered. If the user intends to use a paired primer lying outside the sequence supplied to the program, the user should check that either no exact restriction digests are present in the region not shown to the program or any exact restriction sites will still permit discrimination between diagnostic bands when analyzed with gel electrophoresis. Primers can be chosen based on a strict primer length or by a target melting temperature. A target size may be desired if the user wishes to ensure that a sufficiently large fragment will be cleaved. This may be useful in GC rich areas where a primer designed to match a target melting temperature would be short, resulting in small shifts in band sizes after cleavage and electrophoresis. Melting temperature calculations are performed using the Nearest Neighbor method [10] using thermodynamic parameters published by Sugimoto et al. [11], as implemented in the Oligo Calc tool [12]. It is necessary to assume certain information about the primer concentration and sodium ion concentration in the PCR reaction to calculate the melting temperature. Default values have been provided, but those parameters can be modified as necessary by the user through the web interface. Also, primers which contain terminal 3’ mismatches are rejected by default. Some researchers have reported that certain terminal 3’ mismatches are compatible with PCR [13-15], but due to inconsistencies in the literature, the default assumption for this tool is that 3’ mismatches will not amplify. If the user wishes to allow certain 3’ mismatches, the option is available. If enabled, 3’ G/T mismatches are ignored.

The second box presented by the tool to the user permits screening for mutagenesis events in a CRISPR-mediated mutagenesis experiment. The box requires the user to submit the wild-type genomic sequence the user intends to target. The sequence should contain at least twenty bases on each side of the cleavage site. The user should also include the 5’ to 3’ CRISPR target site, not including the PAM. The CRISPR target sequence is not required to be in the same orientation as the wild-type sequence. The tool assumes that the last base of the provided sequence immediately precedes the PAM if aligned to the wild-type sequence and that cleavage occurs at the -3 position. The mismatch max parameter behaves as it does in the first box. The final major parameter is the acceptable loss threshold, which is the percent of editing events the user is willing to miss with their screening. Lower values mean the user wants to detect more editing events. Higher values mean the user is willing to accept missing certain editing events. Missed editing events, in this context, are most likely to occur if an insertion event occurs relative to an enzyme with degenerate bases in its recognition motif. The advanced options are the same as for those in the first box.

An additional application is facilitated by the first box, the known-alleles tool. CRIPSR-mediated mutagenesis events create random mutations at the target locus. It may be desired in some cases to generate an isogenic mutation in a novel biological context, such as a different ecotype or genetic background. This is especially useful in cases where multiple mutant loci must be maintained and introducing an isogenic mutation would prove easier than screening multiple segregating loci as a result of a cross. The output of the known-alleles tool indicates which of the two supplied sequences is cleaved by the restriction digest. If the user supplies the wild-type sequence and the specific mutant sequence they wish to find and choose an assay where the mutant sequence is cleaved, then a CAPS/dCAPS assay can be used to screen a pool of CRISPR-generated mutants for a specific mutation. This could be feasibly accomplished by a two-step process, where primers generated by the unknown allele indCAPS application are used to screen for lines showing any evidence of CRISPR-mediated editing events, and then a second primer set is used to screen for a specific mutation within that population.

### Technical details of the indCAPS package

A general outline of the algorithm that was developed is illustrated in figure 2. The user-supplied sequences (based on the mutagenesis target) are compared by defining shared and unshared regions in each sequence. In the case of a SNP allele, each sequence will have an unshared region of one base. For indel alleles, the sequence with the deletion relative to the other will have an unshared region of 0 bases and the sequence with the relative insertion will have an unshared region equal to the number of inserted bases.

**Fig 2.**
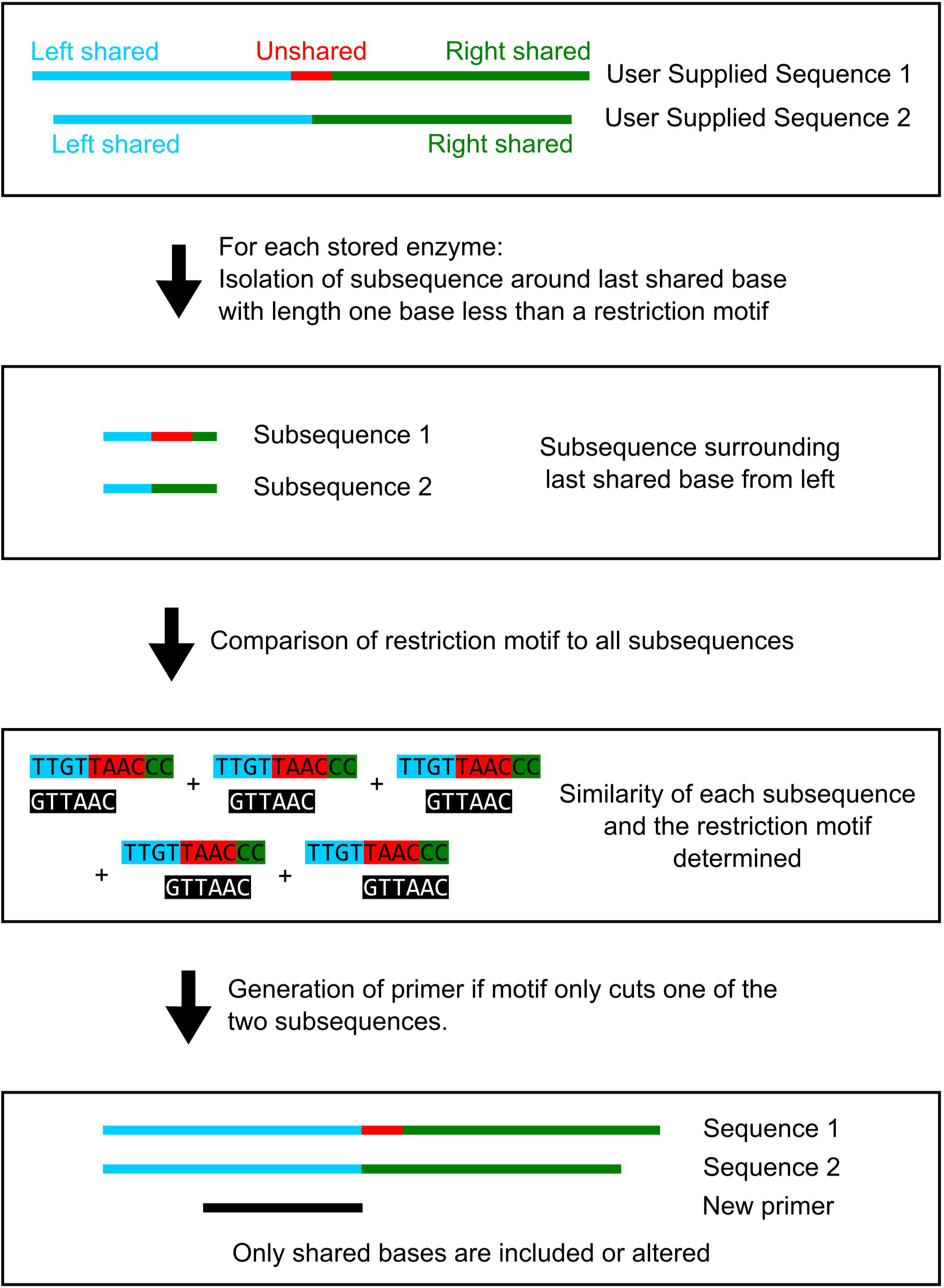
Algorithm for generation of oligonucleotide primers useful for CAPS and dCAPS assays. Two user-supplied sequences are analyzed, with one end near the predicted mutation site. Shared and unshared regions are identified in each sequence. A sub-sequence near the last shared base from each direction is isolated and compared to a library of restriction enzyme recognition motifs (https://www.neb.com/tools-and-resources/selection-charts/alphabetized-list-of-recognition-specificities). If a diagnostic site is detected, determined by an exact motif match in only one sequence, a primer is generated. The primer disrupts any exact matches present in the shared regions and is checked to ensure the mismatch number is less than the specified maximum.

Two core assumptions are made when designing diagnostic assays: 1) designed primers must be wholly contained in the shared region; and 2) putative restriction sites must have at least one base pair of overlap with both the shared and unshared regions. A library of restriction enzyme recognition sites (https://www.neb.com/tools-and-resources/selection-charts/alphabetized-list-of-recognition-specificities) (excluding nicking or double-cutting enzymes) is compared to both shared regions and to a subsequence near the last shared base. An enzyme is rejected if it cuts in the downstream shared region, and exact matches in the primer are disrupted with mismatches during the primer design stage. Each set of sequences is analyzed twice, once as supplied, and again using the reverse complement of each sequence. An enzyme that is rejected because it cuts in the shared, downstream region of both sequences from one direction may be suitable if the primer is aligned to the reverse complement of the two sequences.

For the purpose of CRISPR/Cas9 screening, a profile of possible editing events is used to simulate editing events in the wild-type sequence. Currently, the default editing events are single base pair insertion and deletion events. Future versions of the software will allow the user to supply a custom profile of events. The user is required to provide the specific target sequence used in CRISPR/Cas9 mutagenesis and the cut site is assumed to be at the -3 position of the provided 20 bp target [9]. All possible sequence variants are created and compared to the wild-type sequence to identify the last shared base. The last shared base is taken to be the last base in the wild-type sequence shared with all sequence variants. A simplifying assumption is made when designing primers for unknown alleles: cleavage will occur in the wild-type sequence and will be disrupted in mutant sequences.

### Use of indCAPS to find mutants in cytokinin signaling

Cytokinins, a class of adenine-derived signaling molecules are involved in regulating a diverse set of biological processes. Cytokinins are perceived by Arabidopsis Histidine Kinase (AHK) proteins [16,17] in the endoplasmic reticulum [18,19], which undergo autophosphorylation on a His residue. This phosphate is ultimately transferred to either type-A or type-B Response Regulator proteins (ARRs) via the Histidine Phosphotransfer (AHP) proteins [20-22]. Type-B ARRs are transcription factors activated by phosphorylation [23]. Type-A ARRs lack a DNA-binding domain, are cytokinin-inducible, and negatively regulate cytokinin signaling [24,25].

The cytokinin AHK receptors are encoded by *AHK2*, *AHK3*, and *AHK4/CRE1* in Arabidopsis. Mutant lines with loss-of-function T-DNA insertion alleles of each gene have been identified. The most severely affected triple mutant line,, *ahk2-7 ahk3-3 cre1-12*, harbors null alleles for *ahk2* and *ahk4*, but still contains residual full-length wild-type transcript for *AHK3* [3]. We sought to identify a CRISPR-induced null allele of *ahk3* in an *ahk2 ahk4* background by introducing a frameshift mutation in *AHK3*. This would reveal the effect of complete disruption of the AHK cytokinin receptors in Arabidopsis.

The indCAPS tool was tested by designing primers for a CRISPR mutagenesis experiment targeting the *AHK3* gene (Fig. 2). A CRISPR/Cas9 target was designed to target the first exon of *AHK3* using the CRISPR-Plant resource [8]. The target site chosen is before the Cyclases/Histidine kinases Associated Sensory Extracellular (CHASE) domain, the cytokinin binding domain of the AHKs. The *AHK3* targeting plasmid, pCH59, was stably transformed into an *ahk2-7 cre1-12* mutant line [16,17,26], referred to as *ahk2,4* hereafter. The pCH59 vector was constructed by cloning a commercially synthesized gRNA fragment into a pCUT binary vector system expressing plant codon optimized Cas9 [4]. T1 transformed seedlings were identified by hygromycin selection and grown to maturity. The T2 progeny were screened for *AHK3* editing events using primers generated by indCAPS. Viable seedlings with homozygous, single base pair insertions causing frameshift mutations disrupting the *AHK3* coding region were identified. These alleles, denoted *ahk3-9* and *ahk3-10*, are single base pair insertions of A and C, respectively. The frameshift produces an early stop 25 residues after the edit location. The resulting predicted protein retains two transmembrane domains, but no functional CHASE domain or cytosolic histidine kinase or receiver domains. These triple *cre1-12 ahk2-7 ahk3* mutants were viable and resembled the *cre1-12 ahk2-7 ahk3-3* [27]. These results demonstrate that the complete disruption of all three AHK cytokinin receptors does not result in embryo lethality. As these are the only CHASE-domain containing proteins in Arabidopsis, this suggests that either cytokinin is not essential for early development, or that there are other as yet unidentified cytokinin receptors.

**Fig 3.**
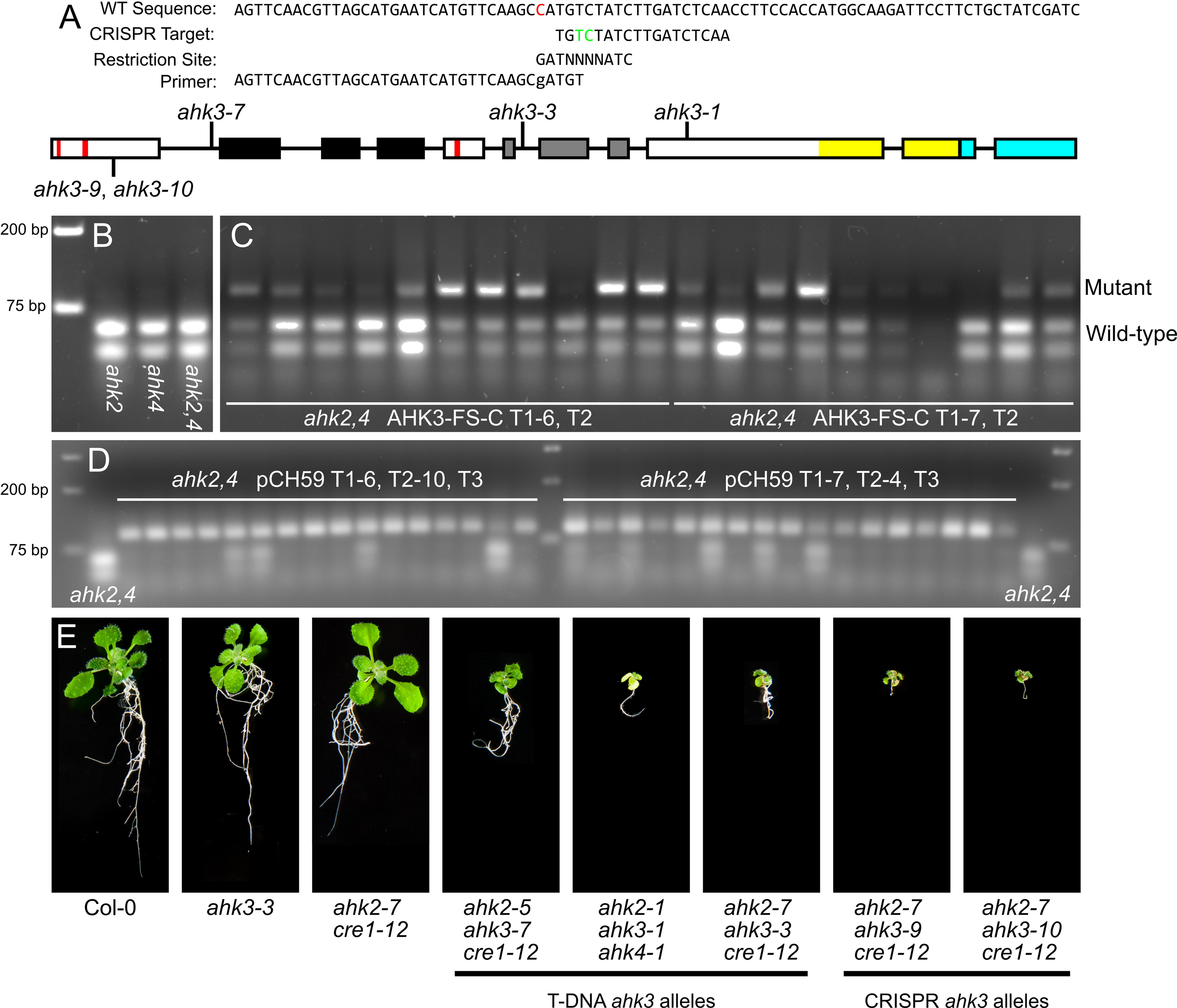
Homozygous editing events in *AHK3* were identified. (A) The indCAPS package was used to generate a primer recognizing a Bsa BI site spanning the CRISPR cut site (between the green bases). A single mismatch was required in the primer (indicated in red). The genomic locus for *AHK3* is shown. Boxes indicate exons, red bars - transmembrane domains, black region - CHASE domain, grey region – histidine kinase domain, yellow region - receiver domain, blue region – 3’ UTR. Locations of T-DNA insertion sites and targeted editing site are indicated. (B,C) The assay was used to screen *A. thaliana* plants stably transformed with a pCUT binary vector system [4] and sgRNA constructs targeting *AHK3*. (B) Wild-type controls at edit location. (C) T2 plants from two representative independent transformation events are shown. Uncut amplicon is 90 bp, wild-type allele is cleaved to produce 36 bp and 54 bp fragments. (D) Offspring from two T2 plants heterozygous for editing events were selected and analyzed for editing. (E) *A. thaliana* seedlings imaged at 2.5 weeks of growth. Shown are Col-0; *ahk3-3; ahk2,4*; *ahk2-5 ahk3-7 cre1-12*; *ahk2-1/+ ahk3-1 ahk4-1*; *ahk2-7 ahk3-3/+ cre1-12; ahk2-7 ahk3-9 cre1-12*; *ahk2-7 ahk3-10 cre1-12*. All plants at same scale.

## Conclusions

The indCAPS package provides a useful tool for researchers using CRISPR-mediated mutagenesis as it facilitates the screening of individuals in which editing of the target has occurred. It also provides replacement for existing tools for the design of primers for dCAPS analysis capable of distinguishing known indel alleles. We employed this tool to successfully design diagnostic primers to identify CRISPR-induced *ahk3* null alleles, the subsequent analysis of which showed that the cytokinin AHK receptors are not essential for embryo development.

## Acknowledgements

We would like to thank G. Eric Schaller, Christian Burr, Amala John, Joanna Polko, Lauren Ross, and Ariel Aldrette for helping with the visual design of the website and the figures for this paper and Jamie Winshell for technical support. This work was supported by a grant from the National Science Foundation (IOS-1238051).

